# Characterizing and evaluating cell specialization through Gini index of gene expression: TCGA normal vs tumor case study

**DOI:** 10.1101/2024.06.19.599717

**Authors:** Fabio Cumbo, Daniele Santoni

**Affiliations:** Center for Computational Life Sciences, Lerner Research Institute, Cleveland Clinic, 9500 Euclid Avenue, Cleveland, 44195 OH, USA; Institute for Systems Analysis and Computer Science “Antonio Ruberti”, National Research Council of Italy, Via dei Taurini 19, Rome, 00185 RM, Italy

## Abstract

In this work we introduce the Gini Index, commonly used as a measure of statistical dispersion to evaluate the income inequality within a nation, as an effective and reliable measure of cell specialization. In particular we use it to evaluate and compare the specialization level of normal and tumor cells according to their gene expressions. Obtained results reveal how Gini Index is able to capture information associated with cell specialization, and show that tumor cells, on average, tend to lose their specialization or in other words their capacity to be the cells they were committed to be, due to cancer.

## BACKGROUND

Gini index (GI) was introduced by the Italian statistician and demographer *Corrado Gini* in the first decades of the 1900s [1,2]. It has been commonly used as a measure of statistical dispersion to evaluate the income inequality within a nation. The general principle is based on the comparison between the portion of economic resources and the portion of the population that possesses those resources. In other words, GI measures in a peculiar way how a distribution of any source of data is far from being a uniform one, collapsing this information in a number ranging from 0 to 1. Although GI is a powerful and effective measure to characterize any sample distribution, it was applied only a few times in a biological research context. Jiang and colleagues developed *GiniClust*, a tool that uses GI in a biological context to characterize rare cell types in single-cell experiments [3]. Tsoucas and Yuan developed a new tool *GiniClust2* that improved the ability to detect and cluster different cell types in single-cell experiments [4]. GI was also used to characterize and identify gene classes according to their expression variability across different cells [5], or to select genes for normalizing expression profiling data [6].

In this work we introduce the GI as an effective and reliable measurement of the specialization of cells, using it to evaluate and compare the specialization level of normal and tumor cells according to their gene expressions.

## RESULTS

The main goal of this paper is to study gene expression specialization through GI index comparing cancer and normal cells and reporting different behaviors. We present a global view of GI values associated with samples coming from patients with different cancer types (see Table S1) for both normal and tumor cells. Secondly we analyze and compare for each single patient, normal and tumor GIs, showing through z-score values they are mostly statistically different. Finally, we study and evaluate for each tumor type statistical differences between normal and tumor GI distributions through Wilcoxon tests. We applied paired statistical tests to compare GI distributions of normal and tumor coupled samples. We then consider a broader dataset including all available samples even if not coupled, trivially applying non paired statistical tests.

Table S2 reports a global view of GI values for each tumor type. The first column indicates the tumor type, the second column indicates the number of subjects for which both normal and tumor samples are available. The other columns show other statistical parameters related to GI distributions. GI values are typically distributed around 0.9, the average in normal samples ranges from 0.907 in BRCA to 0.969 in LICH while in tumor samples from 0.906 in LUSC until 0.951 in LICH (see Table S2). Figure 1 shows the comparison between GI distributions of tumor (orange) and normal (blue) cells for four different cancer types (see Supplementary File D1 for GI data of all cancer types): the panels in the green box - on the left side of the figure - (HNSC and LIHC) clearly show higher GI values for normal samples with respect to tumor samples. On the other hand the panel in the red box - right side of the figure - (THCA) shows an opposite behavior with higher values for tumor samples. The central panel in the black box (PRAD) shows an intermediate case where there is no clear prevalence between tumor and normal samples. Most of the tumor types typically show GI values lower in cancer than in normal (see Supplementary Figure SF1 for a complete view).

**Figure 1:**
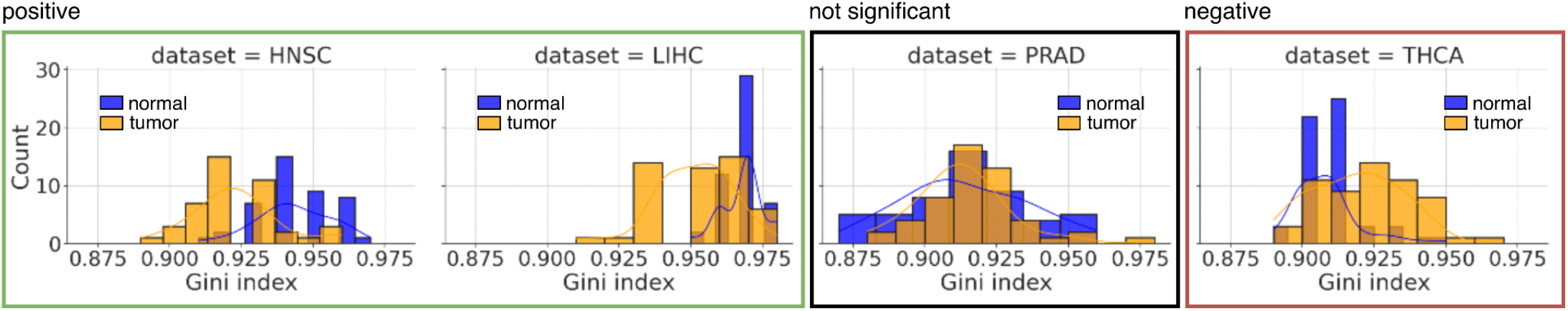
Tumor and normal GI distributions for 4 different tumor types: HNSC and LIHC (green box), THCA (red box) and PRAD (black box). The plots show for each of the 4 considered tumor types how many samples (y-axis), tumor in pink and normal in cyan, have a GI falling in the corresponding bin (x-axis).

In order to statistically evaluate the significance of this difference at a patient level we compare the difference between tumor and normal GIs with the difference distribution between artificial gene expression arrays randomly generated from the actual ones obtaining a z-score value and the corresponding P-value (see Materials and Methods).

In Table 1, the number and percentage of patients showing a significant positive difference between normal and cancer cells are reported in column 3 for each tumor type. In the same way, column 4 and 5 report the numbers and percentages of not significant and significant negative GI differences respectively. The rows corresponding to a given tumor type are highlighted in green (red) when the majority of patients shows a significant positive (negative) difference (P-value < 0.01). In the same Table 1, **+** and **−** symbols depend on the statistical significance of the Wilcoxon rank-sum test performed on all the normal and tumor samples regardless of their pairness (see Material and Methods). A **+** symbol in case the comparison of normal vs tumor samples is significant considering an an adjusted p-value<0.01 according to the Bonferroni correction, and a **−** symbol in case of the tumor vs normal comparison with the same p-value threshold and correction. The two analyses lead to, as expected, similar and consistent results even if they provide a different view at a patient level and at a global level.

**Table 1:** Summary table of the statistical analysis performed on the number of normal-tumor paired samples (second column) for each of the involved 17 tumor types (first column): (**Z-scores**) columns 3–5 report the number of paired samples (and their percentage) for which the p-value, computed considering the z-score of the actual pair and those of the 1,000 Gini indices on the randomized gene expression profiles, is lower than 0.01. In particular, the column “positive” reports the number of significant p-values on positive z-scores, and vice-versa for the “negative” column. Instead, the “not significant” column contains the number of paired samples for which the p-value is not significant, regardless of the positive or negative sign of their z-scores. Results are coded according to the values reported under the 3rd, 4th and 5th columns: green if “positive” sample pairs (3rd column) are the majority (percentage higher than 50%) and red, on the other way around, if “negative” sample pairs (5th column) are the majority (percentage higher than 50%). (**Wilcoxon**) the presence/absence of the and symbols near the tumor type represents the statistical significance according to the Wilcoxon rank-sum test.

| Tumor Type | Samples<br>normal – tumor (paired) | Comparison of Gini indices for each patient through z-score |  |  |
| --- | --- | --- | --- | --- |
| | | Positive<br>($p < 0.01$) | Not significant | Negative<br>($p < 0.01$) |
| BLCA | 19 – 408 (19) | 11 (57.9%) | 6 (31.6%) | 2 (10.5%) |
| BRCA | 113 – 1090 (112) | 34 (30.4%) | 34 (30.4%) | 44 (39.3%) |
| CHOL + | 9 – 36 (9) | 9 (100%) | 0 (0%) | 0 (0%) |
| COAD + | 41 – 456 (41) | 22 (53.7%) | 15 (36.6%) | 4 (9.8%) |
| ESCA + | 11 – 161 (8) | 6 (75.0%) | 2 (25.0%) | 0 (0%) |
| HNSC + | 44 – 500 (43) | 32 (74.4%) | 8 (18.6%) | 3 (7.0%) |
| KICH – | 24 – 65 (23) | 0 (0%) | 1 (4.3%) | 22 (95.7%) |
| KIRC + | 72 – 530 (72) | 35 (48.6%) | 20 (27.8%) | 17 (23.6%) |
| KIRP | 32 – 288 (31) | 9 (29.0%) | 9 (29.0%) | 13 (41.9%) |
| LIHC + | 50 – 371 (50) | 43 (86.0%) | 5 (10.0%) | 2 (4.0%) |
| LUAD | 59 – 513 (57) | 19 (33.3%) | 30 (52.6%) | 8 (14.1%) |
| LUSC + | 49 – 501 (49) | 19 (38.8%) | 23 (46.9%) | 7 (14.3%) |
| PRAD | 52 – 495 (52) | 9 (17.3%) | 27 (51.9%) | 16 (30.8%) |
| READ | 10 – 166 (9) | 1 (11.1%) | 8 (88.9%) | 0 (0%) |
| STAD + | 32 – 375 (27) | 18 (66.7%) | 9 (33.3%) | 0 (0%) |
| THCA – | 58 – 502 (58) | 6 (10.3%) | 20 (34.5%) | 32 (55.2%) |
| UCEC | 35 – 543 (23) | 7 (30.4%) | 5 (21.7%) | 11 (47.8%) |

Wilcoxon tests are also performed on the GIs distributions of paired samples for each tumor obtaining the same results except for LUSC and ESCA that are found not significant.

We always refer to the different tumor types with their abbreviations as reported on the Genomics Data Commons website (see Table S1 for the list of these abbreviations alongside with their extended description).

## CONCLUSIONS

GI is able to characterize a distribution evaluating how close it is from a uniform one. It provides information associated with, but also complementary to, other measures such as the standard deviation or the entropy. However, it has several advantages such as the direct comparability of any set of data since it is a number in the range [0,1]. Moreover, it does not need any kind of assumption as in the case of entropy, where in most cases, a binning supervised pre-process is necessary. In this view it seems particularly suitable to be applied in the context of computational biology. To the best of our knowledge, this work is the first attempt to apply GI to gene expression in the context of tumors comparing the GI of normal and tumor cells. The observed loss of specialization in tumor cells corresponds in our analysis to a lower GI with respect to normal cells. This behavior was observed both at a single patient level comparing GIs of coupled samples (from the same patient) through z-scores analysis and at a global level comparing distributions of GIs in normal and tumor dataset. Interestingly, despite this being the overall typical behavior, few patients show an unexpected increase of their GIs. In the same way not all cancer types display this behavior, some of them show an increase of specialization in tumor cells.

## MATERIALS AND METHODS

We focus on the public gene expression (FPKM - *Fragments Per Kilobase of transcript per Million mapped reads*) quantification experiments of the TCGA program available on the open-access OpenGDC repository [7] for running our analyses. Here, every kind of experimental data and metadata are first extracted from the Genomic Data Commons portal [8].

In this study, we focus on 17 out of 33 different tumor types available in the OpenGDC database considering a number of paired normal-tumor samples ranging from a minimum of 9 (ESCA) to a maximum of 112 (BRCA). The number of samples for each tumor type is reported in Table 1 as well as in Table S2.

For each cancer type and for each patient we compare paired normal and tumor gene expression GIs to assess whether they are significantly different. The actual difference between normal and tumor GIs is compared with a distribution of 1,000 artificial GI differences obtained by randomizing gene expressions of the two samples (see Section Supplementary Methods SM for details).

At the end of this procedure we obtain a z-score associated with each considered patient assessing how many standard deviations the actual difference between normal and tumor deviates from the mean of the randomized GI difference distribution. A positive z-score is associated with a positive difference so that normal GI is significantly greater than tumor GI. In this case normal cells are more specialized than tumor ones. On the other hand a negative z-score is associated with a negative difference so that normal GI is significantly smaller than tumor GI. The adjusted p-values associated with each z-score are then computed and a threshold of 0.01 is set to assess the significance of the difference between normal and tumor GIs.

For each cancer type we also compare paired normal and tumor gene expression GI distributions. To assess whether they come from the same hypothetical distribution we perform paired Wilcoxon tests. Bonferroni adjustment is applied for multiple test corrections. We also compare normal and tumor GI distributions for all available samples taking into account also not paired ones (samples for which only one condition is available). Unpaired Wilcoxon tests and Bonferroni adjustment are applied in the same way.

## Acknowledgement

Daniele Santoni is a member of the National Group for Scientific Computing (GNCS) – Istituto di Alta Matematica Francesco Severi (INdAM).

