## Supplementary Material for "Characterizing and evaluating cell specialization through Gini index of gene expression: TCGA normal vs tumor case study"

#### STUDY ABBREVIATIONS

The 17 different tumor types considered in this study in the form of study abbreviations are all reported in Table S1 alongside with their extended name as reported on the official Genomics Data Commons web portal at https://gdc.cancer.gov/resources-tcga-users/tcga-code-tables/tcga-study-abbreviations.

| Study Abbreviation | Study Name |
|---|---|
| BLCA | Bladder Urothelial Carcinoma |
| BRCA | Breast Invasive Carcinoma |
| CHOL | Cholangiocarcinoma |
| COAD | Colon Adenocarcinoma |
| ESCA | Esophageal Carcinoma |
| HNSC | Head and Neck Squamous Cell Carcinoma |
| KICH | Kidney Chromophobe |
| KIRC | Kidney Renal Clear Cell Carcinoma |
| KIRP | Kidney Renal Papillary Cell Carcinoma |
| LIHC | Liver Hepatocellular Carcinoma |
| LUAD | Lung Adenocarcinoma |
| LUSC | Lung Squamous Cell Carcinoma |
| PRAD | Prostate Adenocarcinoma |
| READ | Rectum Adenocarcinoma |
| STAD | Stomach Adenocarcinoma |
| THCA | Thyroid Carcinoma |
| UCEC | Uterine Corpus Endometrial Carcinoma |

**Table S1:** List of study abbreviations with their extended study names for the 17 tumor types from the TCGA program considered in our analysis as reported on the Genomics Data Commons portal.

#### SUMMARY OF GINI INDICES

Here we reported a summary table of the Gini indices (Table S2) for all the normal-tumor paired samples in each of the tumor types involved in this study, including their average values with the standard deviations and their numerical range (minimum and maximum values).

| Tumor Type | Subjects | Summary of Gini indices | | | |
|---|---|---|---|---|---|
| | | Normal (average ± stdev) | Tumor (average ± stdev) | Normal (min – max) | Tumor (min – max) |
| BLCA | 19 | 0.926 ± 0.010 | 0.916 ± 0.014 | 0.905 – 0.945 | 0.889 – 0.934 |
| BRCA | 112 | 0.906 ± 0.019 | 0.907 ± 0.016 | 0.873 – 0.966 | 0.878 – 0.971 |
| CHOL | 9 | 0.953 ± 0.003 | 0.917 ± 0.015 | 0.964 – 0.964 | 0.902 – 0.953 |
| COAD | 41 | 0.926 ± 0.008 | 0.917 ± 0.013 | 0.910 – 0.944 | 0.892 – 0.962 |
| ESCA | 8 | 0.937 ± 0.016 | 0.912 ± 0.004 | 0.911 – 0.965 | 0.901 – 0.920 |
| HNSC | 43 | 0.943 ± 0.042 | 0.922 ± 0.014 | 0.907 – 0.972 | 0.887 – 0.960 |
| KICH | 23 | 0.908 ± 0.019 | 0.948 ± 0.015 | 0.880 – 0.932 | 0.909 – 0.974 |
| KIRC | 72 | 0.926 ± 0.027 | 0.916 ± 0.015 | 0.891 – 0.955 | 0.878 – 0.949 |
| KIRP | 31 | 0.924 ± 0.026 | 0.925 ± 0.020 | 0.890 – 0.946 | 0.888 – 0.958 |
| LIHC | 50 | 0.968 ± 0.065 | 0.951 ± 0.012 | 0.953 – 0.983 | 0.914 – 0.975 |
| LUAD | 57 | 0.914 ± 0.021 | 0.909 ± 0.013 | 0.893 – 0.939 | 0.870 – 0.939 |
| LUSC | 49 | 0.913 ± 0.021 | 0.905 ± 0.016 | 0.892 – 0.935 | 0.854 – 0.938 |
| PRAD | 52 | 0.913 ± 0.020 | 0.913 ± 0.016 | 0.871 – 0.956 | 0.879 – 0.978 |
| READ | 9 | 0.919 ± 0.024 | 0.918 ± 0.008 | 0.906 – 0.933 | 0.895 – 0.933 |
| STAD | 27 | 0.945 ± 0.043 | 0.924 ± 0.016 | 0.915 – 0.969 | 0.894 – 0.957 |
| THCA | 58 | 0.908 ± 0.019 | 0.918 ± 0.017 | 0.891 – 0.950 | 0.887 – 0.974 |
| UCEC | 23 | 0.911 ± 0.020 | 0.919 ± 0.02 | 0.893 – 0.924 | 0.881 – 0.965 |

**Table S2:** Summary table of the Gini indices of normal and tumor samples for each of the 17 tumor types from the TCGA program (1st column), reporting the number of subject for which both normal and tumor samples are available (paired – 2nd column), the average Gini index with its standard deviation (3rd and 4th columns), followed by the minimum and maximum Gini indices (5th and 6th columns).

### MATERIALS AND METHODS
### Additional material

#### Evaluating differential specialization of normal vs tumor paired samples

For each cancer type and for each patient we compared paired normal and tumor gene expression GIs to assess whether they are significantly different. Given a cancer type we build the set of samples $P = \{p_1, p_2, .. p_n\}$ for which both normal and tumor gene expressions are available. GIs of gene expression associated with each $p_i$ are indicated with $GI_i^H$ and $GI_i^T$ for normal and tumor samples respectively. We define as $GID(p_i) = GI_i^H - GI_i^T$ the difference between normal and tumor GIs. Now we wonder whether $GID(p_i)$ value is statistically significant or in other words whether the observed value can assess that the two conditions show significantly different gene expression distributions when evaluated through GI. $GID(p_i)$ is a pure number that in general depends on the two distributions so we have to compare it with an expected value obtained taking into consideration the two distributions. We generate artificial couples or random gene expression vectors by shuffling the two gene expression ones. For each gene we randomly associate to the first vector one out of the two gene expression values and we assign the other value to the other vector. Once obtained the two randomized vectors we compute the correspondent GIs namely $GI_i^{HR}$

and $GI_i^{TR}$ and finally $GIDR(p_i) = GI_i^{HR} - GI_i^{TR}$. We iterate this procedure 1,000 times obtaining a collection of 1,000 $GIDR(p_i)$ and we then compute the z-score as $Z_i = \frac{GID(p_i) - Average(\,GIDR(p_i)}{Stdv\,(GIDR(p_i)}$ where $Average(GIDR(p_i))$ and $Stdv(GIDR(p_i))$ are the average and the standard deviation of the 1,000 obtained $GIDR(p_i)$ . According to Shapiro-Wilk normality test, performed on sample cases (data not shown), the distribution of $GIDR(p_i)$ can be considered as extracted from a normal distribution so that we are able to compute the P-value from a given z-score. We compute P-values considering separately the two tails of the normal distribution, evaluating p-value for both tails when the normal GI is greater than the tumor one and vice versa when the normal GI is smaller than the tumor one. We set a P-value threshold of 0.01 applying Bonferroni correction with respect to the number of samples of the considered tumor type. At the very end of this procedure we obtain for each tumor type and for each patient a P-value assessing whether the normal and tumor expression values are significantly different from a statistical point of view when evaluated through Gini indexes.
